# Widespread Recombination Suppression Facilitates Plant Sex Chromosome Evolution

**DOI:** 10.1101/2020.02.07.937490

**Authors:** Joanna L. Rifkin, Felix E.G. Beaudry, Zoë Humphries, Baharul I. Choudhury, Spencer C.H. Barrett, Stephen I. Wright

**Author notes:** These authors contributed equally to this work (Joanna L. Rifkin, Felix E.G. Beaudry).

## Abstract

Classical models suggest recombination rates on sex chromosomes evolve in a stepwise manner to localize the inheritance of sexually antagonistic variation in the sex where it is beneficial, thereby lowering rates of recombination between X and Y chromosomes. However, it is also possible that sex chromosome formation occurs in regions with pre-existing recombination suppression. To evaluate these possibilities, we constructed linkage maps and a chromosome-scale genome assembly for the dioecious plant *Rumex hastatulus*, a species with a young neo-sex chromosome found in part of its geographical range. We found that the ancestral sex-linked region is located in a large region characterized by low recombination. Furthermore, comparison between the recombination landscape of the neo-sex chromosome and its autosomal homologue indicates that low recombination rates preceded sex linkage. Our findings suggest that ancestrally low rates of recombination have facilitated the formation and evolution of heteromorphic sex chromosomes.

## 2 Introduction

Plant and animal genomes vary widely in recombination rate, both between and along chromosomes (1), but how this variation contributes to genome evolution remains unclear. Recombination rate evolution is likely to be governed by natural selection because it is predicted to affect local adaptation and speciation (2), the efficacy of selection (3, 4), and the maintenance of genetic polymorphism (5, 6). However, deterministic models suggest that the conditions under which natural selection favors the invasion of variants that change the rate of recombination (recombination modifiers) are quite restrictive (7). Many other factors, including chromosomal position (8), chromatin structure (9) and transposable elements (10) influence rates of recombination, and evolutionary changes in these properties may indirectly drive recombination rate evolution. Nonetheless, compelling evidence supporting the evolution of recombination rate by natural selection is growing (11, 12). Disentangling the relative contributions of direct selection and other factors in shaping the rate of recombination is essential for detailed understanding of the forces shaping genome evolution.

Sex chromosomes are particularly valuable for the study of recombination evolution because they represent an example of convergent recombination suppression, and evolutionary theory predicts an important role for natural selection in this process. Classical models of sex chromosome evolution predict that sex chromosomes evolve from autosomes to alleviate the cost of sexually antagonistic alleles in the sex to which those alleles are deleterious (13, 14). Because of differences between the sexes in their optimal reproductive strategies (15), some alleles beneficial in one sex can be detrimental in the other sex and create a selective load in the population (14, 16). The cost of this genetic load can be resolved by the evolution of sex-specific gene expression (17), or by the invasion of recombination modifiers that link the sexually antagonistic variant with the genomic region responsible for determination of the sex to which that variant is beneficial (18). Thus, sexually antagonistic selection can promote the spread of structural rearrangements including inversions and autosome-sex chromosome fusions (13, 19), or cause the recurrent spread of recombination loss further along the X and Y (20). Theory suggests that this selection has to be very strong, and almost equal and opposite between the sexes (21, 22), for recombination modifiers to invade. Over time, according to these models, recombination suppression spreads in a stepwise fashion along an incipient sex chromosome, leaving a pattern of “strata” with distinct levels of divergence between the X and the Y. Evidence for evolutionary strata between sex chromosomes has been found in several unrelated organisms, including humans (23), chickens (24), and the plant *Silene latifolia* (25). Plant sex chromosomes are generally in earlier stages of divergence than the vertebrate sex chromosomes that led to the development of this “strata” model (26) and are therefore especially useful for identifying the earliest stages of recombination evolution across sex chromosomes.

*Rumex hastatulus* is one of a relatively small number of dioecious plants with heteromorphic sex chromosomes (27). It offers a unique model for the study of sex chromosome evolution because of a polymorphism in sex chromosome karyotype (28). Males to the west of the Mississippi River have four autosomes and an XY pair (XY cytotype), while males to the east of the Mississippi River have three autosomes, a single larger X chromosome and two Y chromosomes (XYY cytotype). Cytological studies (29, 30), as well as patterns of chromosome number and size (28, 31), suggest that this sex chromosome polymorphism arose from a Robertsonian fusion between the ancestral X chromosome and an autosome. The species’ current polymorphic karyotype includes both the ancestral and the derived state of this chromosome. *Rumex hastatulus* therefore provides an unusual opportunity to investigate recombination rates before and after linkage to the sex-determining region. The principal goal of our study was to understand the changes in recombination rate associated with sex chromosome turnover in *R*. *hastatulus* and to relate our results to the classic model of sex chromosome evolution.

Here, we present a chromosome-scale genome assembly of *R. hastatulus* and describe the patterns of recombination and genome content on the sex chromosomes and autosomes. With these data, we investigate whether the pattern of recombination rate heterogeneity and polymorphism is consistent with the classic stepwise model of sex chromosome evolution.

## 3 Results

### 3.1 The sex-linked region in the XY cytotype exhibits low sex-averaged recombination rates

For the genome assembly, we sequenced PACBio reads for one male individual from the XY cytotype (see supplementary materials). Following *de novo* assembly of PACBio SMRTcell reads, contigs were assembled into longer scaffolds using *in vitro* reconstituted chromatin Chicago libraries (32), followed by Hi-C chromosome conformation capture libraries (33, 34). This primary assembly comprised 1.647 Gb of the *R. hastatulus* genome, with half of the genome assembled into 25 scaffolds larger than 11.886Mb (N50). Our assembly size is consistent with C-values for *R. hastatulus* based on flow cytometry (30; Table S1), representing approximately 92% of the estimated total genome size.

Linkage mapping allowed for further scaffolding and correction of misjoins to assemble the genome into the expected five major scaffolds, representing the five chromosome pairs of the XY cytotype of *R. hastatulus* (28). Using RNAseq from 96 individuals representing both sexes (see Table S2 for sex information and supplementary methods for linkage mapping information), we constructed the final genetic map from 988 independent markers on five linkage groups (Table S3). These five chromosomal scaffolds comprised a total of 1.08Gb, or 65% of the primary assembly. Based on the patterns of recombination suppression, our linkage mapping indicated three sub-metacentric and two metacentric chromosomes (green in Figure 1A), which is consistent with the microscopic karyotype (28).

**Figure 1.**
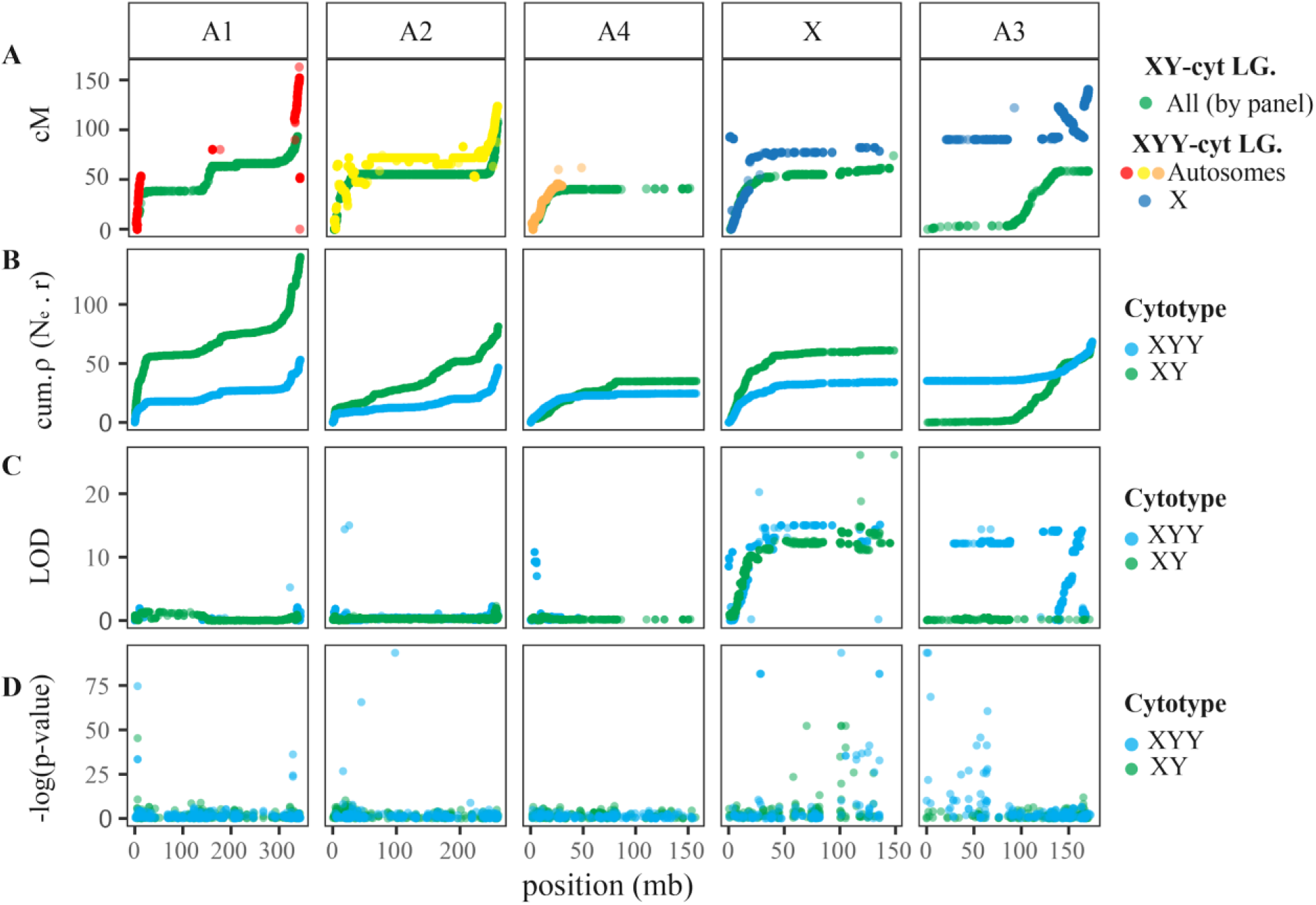
Recombination rate (cM), cumulative recombination rate (ρ), QTL LOD for sex, and GWAS for sex across the five chromosomes of the XY-cytotype *Rumex hastatulus*. **A.** Marey map relating linkage map recombination position (cM) to physical genetic position in the XY-cytotype *R. hastatulus* genome assembly (Mb - panels are XY-cytotype chromosomes) for the XY cytotype (all green) and XYY cytotype (LG split by color). **B.** Cumulative effective recombination rate estimated from population-level genetic sequencing for the XY cytotype (green) and XYY cytotype (blue). **C.** QTL analysis for sex as a binary trait, for the XY cytotype (green) and XYY cytotype (blue). **D.** GWAS for association with sex in population GBS data from (36) for the XY cytotype (green) and XYY cytotype (blue).

Using our chromosome-level assembly, we first used existing RNAseq data from a within-population cross (35) to identify SNPs showing X-linked, Y-linked, X-hemizygous, and autosomal segregation. From this, one linkage group contained the majority of X- and Y-linked SNPs, as well as loci following hemizygous inheritance patterns (Table S4). In particular, of the sex-linked SNPs identified on our five major chromosomal scaffolds, 97% of Y-linked SNPs, 96% of X-linked SNPs and 70% of hemizygous SNPs were mapped onto this scaffold, hereafter referred to as the X. Overall, 52% of sex-linked SNPs were mapped to the major chromosomal scaffolds, indicating that a significant proportion of the sex chromosome sequence could not be positioned in our linkage map, but was present in our assembly as smaller scaffolds. In contrast, 84% of autosomal SNPs were mapped to the major scaffolds; this suggests that the X chromosome assembly is less complete, likely due to the high effective heterozygosity in joint assembly of X and Y chromosomes, and to high repeat density on the Y (30). As expected, windowed analyses of sex-associated SNPs from the experimental cross used for the linkage map showed largely similar patterns. With a quantitative trait locus (QTL) mapping approach, all markers above 9.78Mb on the X chromosome were significantly associated with sex phenotype (LOD, *P*<0.01, Figure 1C, 2D). Thus, overall, we have identified a very large (>140Mb) sex-linked region comprising the vast majority of the X chromosome. The SNP segregation patterns and QTL analyses also suggested the presence of a pseudoautosomal region of X-Y recombination of about 9-10Mb at the beginning of the chromosome (Figures 1 and 2).

Since cross-based analysis may not capture rare recombination events between the X and Y chromosomes, we also used population-level RNA sequence data (35, 36) to assess the boundaries of the sex-linked region. We identified sites as fixed differences between the X and Y chromosome when all males were heterozygous and all females were homozygous, allowing for no genotyping errors. The vast majority of X-Y fixed differences in the population were on the X (Table S5). At the population level, the region of X-Y fixed differences was slightly narrower than in the crossing data, suggesting that this approach did indeed capture more recombination events (Figure 2B). We also conducted a genome-wide association study (GWAS) for sex phenotype using population-level genotyping-by-sequencing (GBS) data (36). We found that regions with significant association with sex, after correcting for multiple hits, were located on the X (Figure 1D, 2E). As these multiple, largely independent approaches converged on one region of one chromosome and showed a significant association with sex, we conclude that we have effectively identified the X-chromosome.

**Figure 2.**
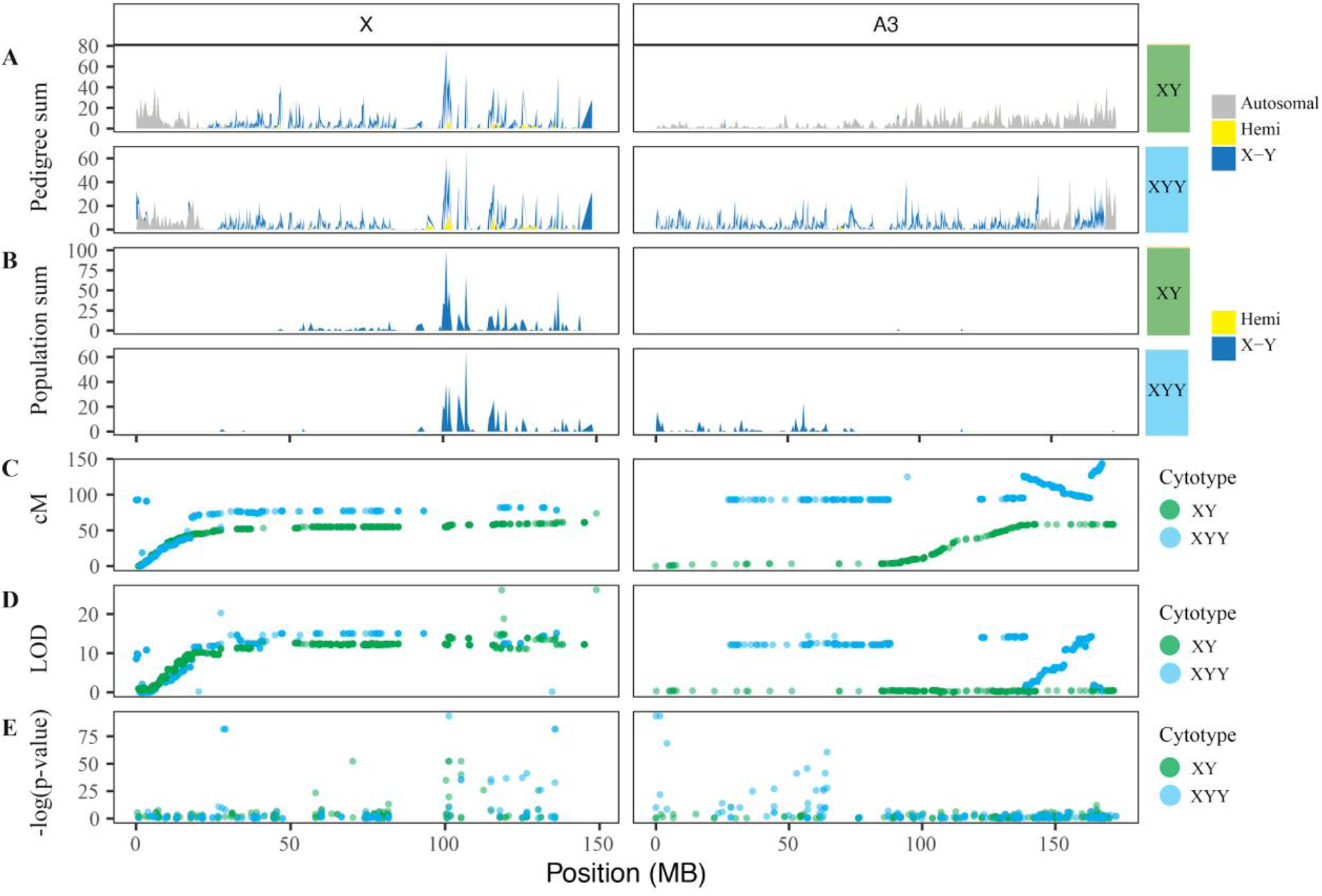
Sex-specific associations across the ancestral X chromosome (panel **X**) and the neo-X chromosome (panel **A3**). **A-B.** Sum of SNPs identified as autosomal (gray – **A** only), hemizygous (yellow), and X- or Y-linked (dark blue) for the XY cytotype (top panel) and XYY cytotype (bottom-panel) in **A.**, cross data from (35), and **B.**, population-wide data from (35, 36). **C.** Marey map relating linkage map recombination position (cM) to physical genetic position in the XY-cytotype *Rumex hastatulus* genome assembly (Mb - panels are XY-cytotype chromosomes) for the XY cytotype (green) and the XYY cytotype (blue). **D.** QTL analysis of sex as a binary trait, for the XY cytotype (green) and XYY cytotype (blue). **E.** GWAS for association with sex in population GBS data from (36) for the XY cytotype (green) and XYY cytotype (blue).

112Mb of the 151Mb X-chromosome were within 9cM of each other. This suggests that the sex-averaged recombination rate in the sex-linked region of the X chromosome was very low (<0.1 cM/Mb). This does not simply reflect suppressed recombination between the X and Y, because our estimate of recombination rate was sex-averaged. Even if X-Y recombination in males was zero, we would estimate a female recombination rate of 0.16 cM/Mb in this region, which is much lower than the average of 1.87 cM/Mb in the pseudoautosomal region. To determine whether this pattern of low recombination was reflected at the population-level scale, we estimated ρ, the effective recombination rate (Ne • r), using LD_hat (37, 38) from our population-level RNAseq sampling data (35, 36). At the population level, which is also sex-averaged, we observed very low effective rates of recombination in the sex-linked region (Figure 1B). Since the low rate of recombination is likely to be the result of X-X as well as X-Y recombination rates, recombination may have been ancestrally low in this region. Low rates of recombination are common in regions surrounding the centromere (39), and it therefore seems likely that the sex-linked region is within a very large pericentromeric region. We observed considerable heterogeneity in the extent of X-Y differentiation along the X chromosome (Figure 2B). Although this is consistent with our previous results, which also identified varying levels of divergence possibly consistent with strata (35), a clear stepwise pattern is not apparent from the ordered data. However, the suppression of recombination limited our ability to order scaffolds based on recombination position in the non-recombining region, and our results could suggest the presence of 2-3 strata on the X chromosome.

### 3.2 Neo-XY shows low recombination rates

To determine whether low rates of recombination are a widespread feature of sex chromosomes in *R. hastatulus*, we also considered the recombination rate on the neo*-*X in the XYY cytotype. We constructed a linkage map using 877 independent markers for the XYY cytotype, complementary to the XY-cytotype map (Figure 1A). Recombination rate estimates indicated that the linkage groups represent three metacentric chromosomes and one submetacentric chromosome, suggesting a reduction in chromosome count, as expected given the X-autosome fusion and consistent with the karyotype of this cytotype (28). A single large metacentric linkage group in the XYY-cytotype linkage map joined the XY-cytotype X chromosome with chromosome A3, suggesting A3 in the XY cytotype is the autosomal homologue to the neo-X chromosome. Our linkage map also identified a large inversion on the recombining end of the neo-X. Inversions have been shown to be important in the evolution of sex chromosomes, as well as in ecological differentiation and reproductive isolation (11, 12). Using cross data from the XYY cytotype from (35), we identified X- and Y-linked SNPs on both the ancestral (panel X) and neo-X (panel A3) sections of the large fused X chromosome (Figure 1B, 2C; Table S4).

Our windowed analyses of sex-linked SNPs also confirmed that sex-linkage extends across both the ancestral X and the neo-X sex chromosomes (Figure 2A, 2B), consistent with an X-autosome fusion event. At the population level, we observed fixed X-Y differences on both the ancestral and neo-X, but the region of fixed X-Y differences on the ancestral X was less extensive than in the XY cytotype (Figure 2B). Both QTL analysis (Figure 2D, LOD P<0.01) and GWAS (Figure 2E, lnL, *P*<0.01) of the XYY cytotype identified large low-recombination regions on both the X and A3 chromosomes to be significantly associated with sex phenotype.

Like the ancestral X chromosome, the sex-linked region of the neo-X exhibited a very large non-recombining region: 107Mb of the neo-X is characterized by a total of 0.56cM (0.0052cM/Mb). However, the severity of recombination suppression is even greater in the context of the large fused chromosome. Interestingly, recombination rates on these chromosomes were similar between the cytotypes, including a 92.16Mb region of 5.682cM (0.074cM/Mb) on A3 (the homologue of the neo-X), despite A3 segregating independently from the sex chromosomes and showing no signal of sex linkage in the XY cytotype (Figure 1A, 2A). This pattern accords with our population-level estimates of recombination rate (Figure 1B). Taken together, this evidence implies that strong recombination suppression in the genomic region that formed the neo-sex chromosome of the XYY-cytotype in *R. hastatulus* preceded its status as a sex chromosome.

### 3.3 Low recombination is a genome-wide phenomenon

The sex chromosomes of *R. hastatulus* were not unusual in exhibiting suppressed recombination. Our analyses revealed that all autosomes also had massive (>100Mb) regions of suppressed recombination, with evidence for recombination restricted primarily to the very tips of the chromosomes (Figure 1A, B). This striking finding is consistent with patterns observed from comparative data, which suggest that species with large chromosomes, such as *R. hastatulus*, often have highly peripheral recombination (8). However, *R. hastatulus* appears to represent an extreme case, with all chromosomes exhibiting over one hundred megabases with recombination rates near zero. Remarkably, approximately 80% of the genome, including the sex chromosomes, exhibits highly suppressed recombination, and given that our linkage map contains only 65% of the complete genome this is likely an underestimate.

## 4 Discussion

The major finding of this study is that the sex-linked regions of both cytotypes of *R. hastatulus* are embedded in a region of highly suppressed recombination which cannot be explained simply by a lack of X-Y recombination. The classic model of sex chromosome evolution assumes that the invasion of recombination modifiers is subsequent to the appearance and maintenance of sexually antagonistic variants (13, 18). However, this may not be necessary if the region already has low rates of recombination (40, 41). In regions with low rates of recombination, the cost of the invasion and maintenance of sexually antagonistic variation can be avoided from the start. Indeed, low rates of recombination are predicted to increase the likelihood of the maintenance of sexually antagonistic polymorphisms (18, 22, 40, 42). Thus, regions of the genome with low rates of recombination may be generally predisposed to evolve sex-linked regions (40). Indeed, evidence for a role for ancestrally low rates of recombination in the evolution of sex chromosomes has started to emerge in plants, as reported in papaya, *Carica papaya*, (43) and kiwifruit, *Actinidia chinensis*, (45). Analogously, self-incompatibility alleles in *Petunia* have been identified in a region of low recombination close to a centromere (45). By comparing a neo-sex chromosome to its ancestral autosome, we provide direct evidence suggesting that suppressed recombination was indeed the ancestral state prior to the evolution of sex-linkage.

This ancestral state is part of a genome-wide global pattern of large, recombination-suppressed chromosomes that produces extremely high linkage disequilibrium even in a dioecious obligate outcrosser. The recombination landscape we observed, of high recombination near chromosome ends and suppressed recombination away from the tips, is consistent with a “telomere-initiate” model of recombination interference (8, 46). Simulations suggest this pattern can lead to heterogeneity in divergence between incipient species at chromosome centers (47), a pattern that may be similar to sex chromosome evolution. Although recombination rates have not yet been quantified in other *Rumex* species, it is noteworthy that sex chromosomes have arisen only in sections of the genus with reduced chromosome numbers (48).

Our results, indicating that low recombination is likely ancestral to the evolution of sex and neo-sex chromosomes in *R. hastatulus*, may seem at odds with two earlier findings from our work on this species: 1) Here we have shown that recombination rate is low in both males and females in this region, yet previous polymorphism analyses reported dramatically lower amounts of genetic diversity caused by linked selection on the Y but not on the X chromosome (49); 2) Given that ancestrally low recombination removes the requirement for the recruitment of recombination modifiers for sex chromosome expansion, how can we also account for our earlier results demonstrating that genes with pollen-biased expression are disproportionately found on the Y chromosome (50)?

The resolution to these apparent contradictions may involve the nonlinear dynamics that likely occur in the transition from low rates of recombination to an absence of recombination. When recombination is low, even very small changes to recombination rate can have important effects on the fixation of adaptive or deleterious mutations (51). For example, simulations suggest that a very low rate of recombination in the European common frog, *Rana temporaria*, can account for the maintenance of high genetic diversity on the Y (52) and, similarly, of high genetic diversity across the genome of the largely asexual apomictic goldilocks buttercup, *Ranunculus auricomus* (53). In our case, a low rate of recombination on the X in females of *R. hastatulus* is not the same genomic environment as the complete absence of recombination on the Y, and the low rates of recombination on the X may be sufficient to maintain genetic diversity in this region. This would imply that the Y and possibly the neo-Y chromosomes have experienced additional recombination suppression.

If this interpretation is correct and recombination suppression has progressed further on the Y, what is the relevance of an initially low-recombination origin for the sex chromosomes? Low but non-zero rates of recombination are in fact likely to be the optimal parameter space for the invasion of recombination modifiers. Indeed, very tight linkage is required for the maintenance of sexually antagonistic polymorphism and, subsequently, for the invasion of recombination modifiers on the sex chromosomes (13). The very low, but non-zero, rates of recombination on the sex chromosomes demonstrated in our study are thus suitable for the maintenance of polymorphism along the proto-sex chromosomes, for example, see Figure 1 in (22). Low rates of recombination may allow for the invasion of recombination modifiers completely linking haploid-expressed (pollen) genes to the Y (50, 54). Therefore, there are theoretical reasons to expect that pre-existing low-recombination plays a crucial role in sex-chromosome formation (40), and in both kiwifruit and papaya, sex-determining regions originated in centromeric regions that were likely ancestrally recombination-suppressed (43, 44). In both of these systems, however, the sex chromosomes are small and homomorphic or micro-heteromorphic. In *R. hastatulus*, the massive genome-wide scale of recombination suppression may have contributed to the ongoing formation of large, heteromorphic sex chromosomes. The ancestral recombination landscape may thus be a major determinant of the genomic structure of sex chromosomes.

## 5 Methods

### 5.1 Preliminary genome assembly

We sent three grams of leaf tissue from an *R. hastatulus* male F_1_ from two parents from Wesley Chapel, TX (TX-WES in (55)) to Dovetail Genomics LLC, Santa Cruz, California 95060, USA. At Dovetail, high molecular weight (HMW) DNA was extracted and sequenced on 15 PacBio SMRTcells (Single Molecule, Real-Time; (56)). The sample was sequenced to 35x coverage for a total of 6.7M reads. After error correction, 5.7M reads were retained (24x coverage) with an N50 of 9.5kb. The error-corrected reads were assembled by Dovetail Genomics into a primary assembly using Falcon (57, 58) and the assembly was polished with Arrow from the PacBio GenomicConsensus toolkit (https://github.com/PacificBiosciences/GenomicConsensus). This assembly yielded 43,461 contigs with an N50 of 74.7kb.

We sent an additional three grams of leaf tissue from a full-sib male for two orders of further scaffolding by Dovetail Genomics to improve the primary PacBio-Falcon assembly. The assembly was first scaffolded using the Chicago technique (32), which uses *in vitro* reconstituted chromatin for positioning. Two Chicago libraries were sequenced from ~500ng of HMW gDNA reconstituted into chromatin *in vitro* and fixed with formaldehyde. Fixed chromatin was digested with DpnII, 5’ overhangs were filled in with biotinylated nucleotides, and free blunt ends were ligated. After ligation, crosslinks were reversed and the DNA purified from protein. Purified DNA was treated to remove biotin that was not internal to ligated fragments. The DNA was sheared to ~350bp mean fragment size and sequencing libraries were generated using NEBNext Ultra enzymes and Illumina-compatible adapters. Dovetail isolated biotin-containing fragments using streptavidin beads before PCR enrichment of each library. Libraries were sequenced on an Illumina HiSeq X. The number and length of read pairs produced was: 189 million, 2×150bp for library 1; 170 million, 2×150bp for library 2. Together, these Chicago library reads provided 33.42x physical coverage of the genome (1-100kb).

These reads were used to scaffold the PacBio-Falcon assembly using the HiRise pipeline (32) which is designed specifically for using proximity ligation data to scaffold genome assemblies. Dovetail conducted an iterative analysis. They aligned Shotgun and Chicago library sequences to the draft input assembly using a modified SNAP read mapper (http://snap.cs.berkeley.edu). The separations of Chicago read pairs mapped within draft scaffolds were then analyzed by HiRise to produce a likelihood model for genomic distance between read pairs, and the model was then used to identify and break putative misjoins, to score prospective joins, and to make joins above a threshold. The longest scaffold increased from 600kb to 1,977kb, and the L50/N50 increased from 0.075Mb in 6,213 scaffolds to 0.248Mb in 1,887 scaffolds.

We further improved the Chicago-scaffolded assembly using 2 Hi-C chromosome conformation capture libraries from Dovetail (33, 34). Briefly, for each library, chromatin was fixed in place with formaldehyde in the nucleus and extracted. Fixed chromatin was digested with DpnII, 5’ overhangs were filled in with biotinylated nucleotides, and free blunt ends were ligated. After ligation, crosslinks were reversed and the DNA was purified from protein. Purified DNA was treated to remove biotin not internal to ligated fragments. The DNA was then sheared to ~350bp mean fragment size and sequencing libraries were generated using NEBNext Ultra enzymes and Illumina-compatible adapters. Streptavidin beads were used to isolate biotin-containing fragments before PCR enrichment of each library. Dovetail sequenced these libraries on an Illumina HiSeq X. The number and length of read pairs produced for each library was: 158 million, 2×150bp for library 1; 194 million, 2×150bp for library 2. Together, these Dovetail HiC library reads provided 87.47x physical coverage of the genome (10-10,000kb). After assembly with HiRise, the N50/L50 of the assembly increased to 11.89Mb in 25 scaffolds and a longest scaffold of 146,334kb.

We obtained C-values for genome size estimation from Plant Cytometry Services of Didam, The Netherlands. We shipped fresh leaf tissue, and DNA content was estimated with flow cytometry relative to *Vinca minor* with both DAPI and PI staining.

### 5.2 Linkage mapping

We generated F_2_ linkage-mapping populations for both the XY and XYY cytotypes of 96 offspring each. The original parents were collected from Wesley Chapel, TX and Marion, SC, respectively (see (55)). Seeds from F_1_ plants were sterilized in 5% (V/V) bleach for one minute and then washed in running tap water and distilled water. Sterilized seeds were spread on wet filter paper in Petri dishes and incubated in the dark at 4°C to germinate. After germination (usually within 2-3 weeks), we transplanted seedlings into 6-inch plastic pots filled with Promix^®^ soil and sand (3:1 ratio) with 300ml of Nutricote^®^ (14:13:13, slow releasing fertilizer) for each 60lbs of Promix soil. We grew seedlings in a glasshouse set for 22°C daytime, 18°C nighttime temperature and 16-hour day length at the University of Toronto St. George campus. We watered on alternate days and randomized pots twice weekly for uniform growing conditions and to avoid edge effects. We phenotyped plants for sex at onset of flowering. After plants were sexed, we collected approximately 30mg of young and healthy leaf tissue from individual plants and flash froze it in liquid nitrogen for RNA extraction. Total RNA was extracted using Spectrum™ plant total RNA Kit (Sigma-Aldrich) according to manufacturer’s instructions. We sent RNA samples to Genome Quebec Innovation Centre (McGill University), Montreal, QC for library preparation and sequencing. Libraries were prepared using NEB® mRNA stranded library preparation method and sequenced on two lanes of Illumina NovaSeq S2 PE100 (2×100) sequencing platform using 96 barcodes. A total of ~6.25 billion reads (6,244,145,277) were generated, ranging from ~13 to ~104 million reads per sample with an average of ~32.5 (32,521,589) (median ~28 million (28,041,358).

We aligned samples to the *R. hastatulus* TX draft assembly using STAR 2-pass (STAR_2.5.3a (59)). We sorted alignments, assigned read groups, and marked PCR duplicates using PicardTools 2.18.21-SNAPSHOT [https://broadinstitute.github.io/picard/]. We called variants with bcftools 1.9-67-g626e46b mpileup and call, and filtered using bcftools view for minimum sample depth 10, minimum sample quality 10, minimum site quality 50, allele frequency between 0.1 and 0.9, and minimum 25 individuals called. For use in linkage mapping, we further filtered our variants using bcftools filter and vcftools (60) to remove mislabeled individuals and sites with more than 5% missing data. For linkage mapping, we converted data from vcf to 012 format using vcftools and transformed to csvr format using custom scripts.

We generated a linkage map using the R package ASMAP (61), which implements the minimum spanning tree algorithm in an R/qtl-compatible interface (62, 63). We removed two and three individuals from the XY and XYY mapping populations, respectively, because of apparent contamination. Individuals with high missing data and genetic clones were removed. We generated our XY linkage map using only markers on the 100 biggest scaffolds of our draft assembly and 322 smaller scaffolds that we identified as sex-linked, which contain 1.2Gb of sequence data. Our XYY linkage map used only markers on the 100 biggest scaffolds of our draft assembly. We filtered our initial set of variants (25010 XY / 25544 XYY) to remove colocalized markers (leaving 2255 XY / 2103 XYY) and distorted markers. We constructed the final maps from 988 and 877 markers for XY and XYY, respectively (see Table S1 for progeny sample sizes). We used custom scripts to fuse colocalized markers into the map to maximize the number of scaffolds in the chromosome-scale assembly.

For the XY cytotype, we recovered five major linkage groups with over 100 markers each (108-276), as well as six minor fragmentary linkage groups (5-36 markers). The sizes of the major linkage groups ranged from 75.52 to 108.91 cM, with a total map length of 550.126. This is broadly consistent with expectations for a linkage map with five chromosomes. Recombination frequency (cM) roughly correlates with physical length (bp) (Figure S4). We recovered four major linkage groups with over 100 markers each (103-258) for the XYY cytotype, as well as seven minor fragmentary linkage groups (2-52 markers). The largest minor linkage group, LG11, was largely collinear with the sex chromosome (LG10), containing the same scaffolds and overlapping positions. The sizes of the major linkage groups ranged from 61.74 to 167.07 cM, with a total map length of 584.819 cM. We assessed recombination rates visually using Marey maps (64) relating physical position of markers along scaffolds to recombination position along chromosomes.

We used Chromonomer (http://catchenlab.life.illinois.edu/chromonomer/) to relate our linkage map to our genome assembly. We conducted manual edits to our linkage map before using Chromonomer, to position scaffolds unique to minor linkage groups on major linkage groups based on physically nearby markers and to remove minor linkage groups. We used a script from Nucleomics-VIB’s BioNano Tools (https://github.com/Nucleomics-VIB/bionano-tools) to convert Dovetail’s assembly table file into an .agp file. With Chromonomer, we were able to place 1.09GB (65%) of the Dovetail assembly into five main pseudomolecules representing the major chromosomes. For downstream analyses, we created custom Python scripts using the .agp file output from Chromonomer to translate scaffold positions from our draft assembly to positions along the chromosomes.

### 5.3 Sex-linked variant calling and filtration

For RNA samples from (35, 36), we aligned samples to the *R. hastatulus* XY draft assembly using STAR 2-pass STAR_2.5.3a (59). We aligned reads for GBS samples from (65) using NextGenMap (66). For both alignments, we sorted the reads and assigned read groups using PicardTools 2.18.21-SNAPSHOT [https://broadinstitute.github.io/picard/]. We marked PCR duplicates for RNASeq but not for GBS data. For both datasets, variants were called using bcftools 1.9-67-g626e46b mpileup as described above. We filtered the RNAseq datasets for minimum sample quality 20, minimum site quality 20, minor allele frequency greater than 0.04, and no missing data. We filtered the GBS dataset for minimum sample quality 20, minimum site quality 10, minimum mean depth of 6, minor allele frequency greater than 0.05, and no more than 50% missing data. We converted data for windowed analyses from vcf to 012 format using vcftools.

We identified SNPs as X-linked, Y-linked, hemizygous, autosomal, or male-only expressed in the cross data, the F_2_ data, and the population data using the 012 files described above and custom R scripts based on the segregation patterns described in (35). We converted all sites to positions along chromosomes using custom Python scripts and the Chromonomer .agp file. We summed the different categories of sites across 500kb windows using custom R scripts.

### 5.4 Quantitative trait locus (QTL) mapping and genome-wide association study (GWAS)

We performed QTL mapping of sex as a binary phenotype using the scan1 function of R/qtl2 (67) using the F_2_ mapping population. We adjusted eta_max (the maximum value for the linear predictor in the model) downwards until the model was able to converge. We performed a permutation analysis to identify significance thresholds. We performed GWAS analysis in Gemma (68) using a likelihood ratio test for significance.

## Supplementary Materials

## Supplementary Tables

**Table S1.** C-values were calculated from two XY-cytotype individuals [an F3 Male (ID: TxTx P5 MG) and F3 Female (ID: TxTx P5 MG)] of *Rumex hastatulus*

|  |  |
| --- | --- |
|  | XY |
| Female | 1.8 |
| Male | 1.99 |

**Table S2.**
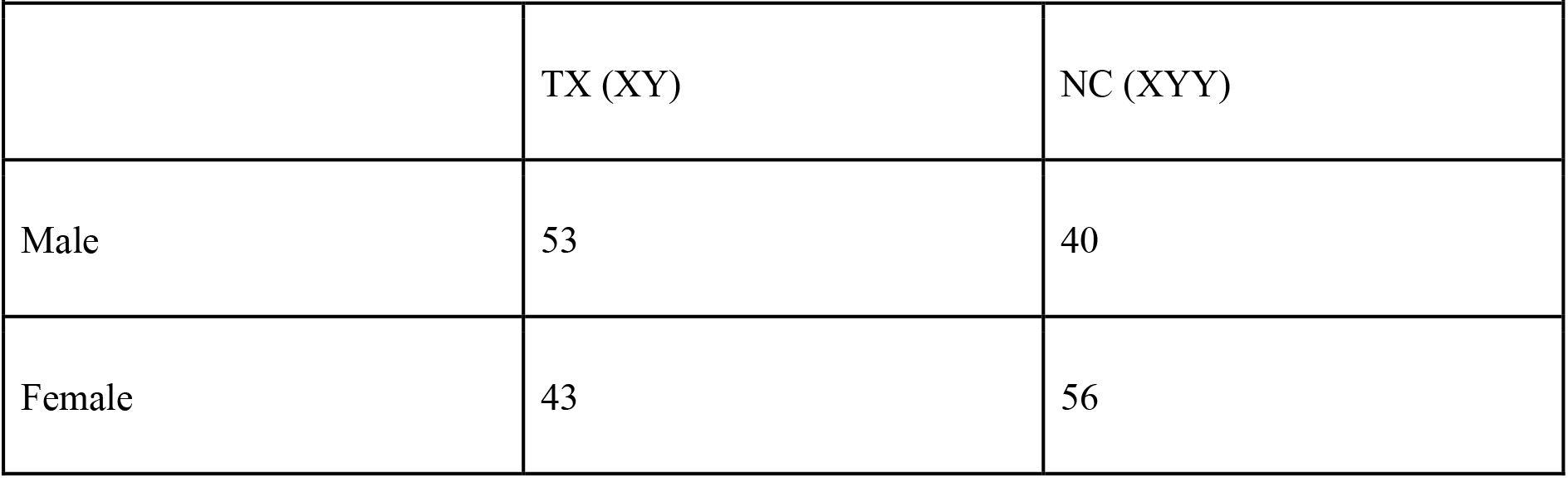
Counts of *Rumex hastatulus* individuals of each sex and cytotype sequenced for the linkage map.

|  |  |  |
| --- | --- | --- |
|  | TX (XY) | NC (XYY) |
| Male | 53 | 40 |
| Female | 43 | 56 |

**Table S3.**
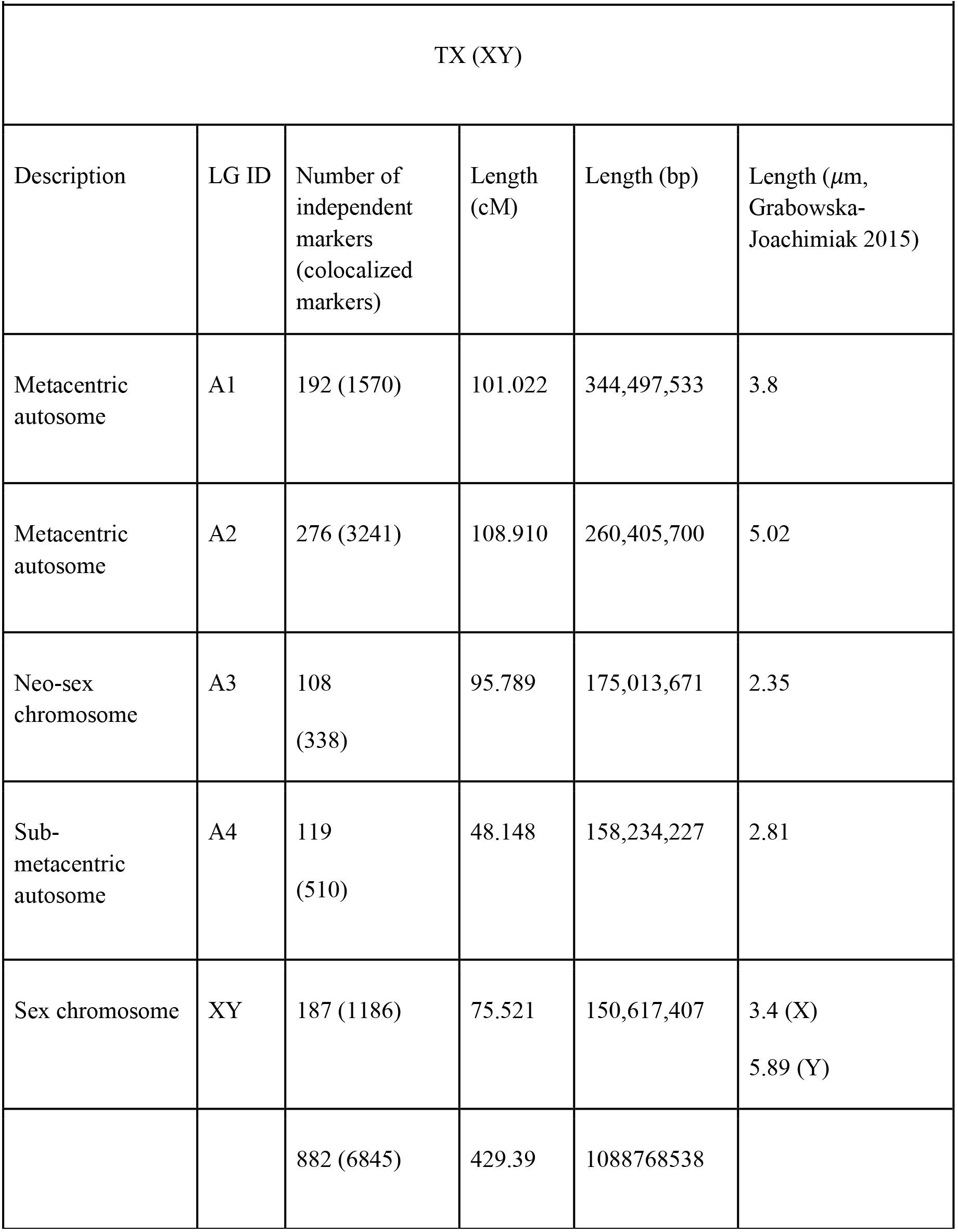
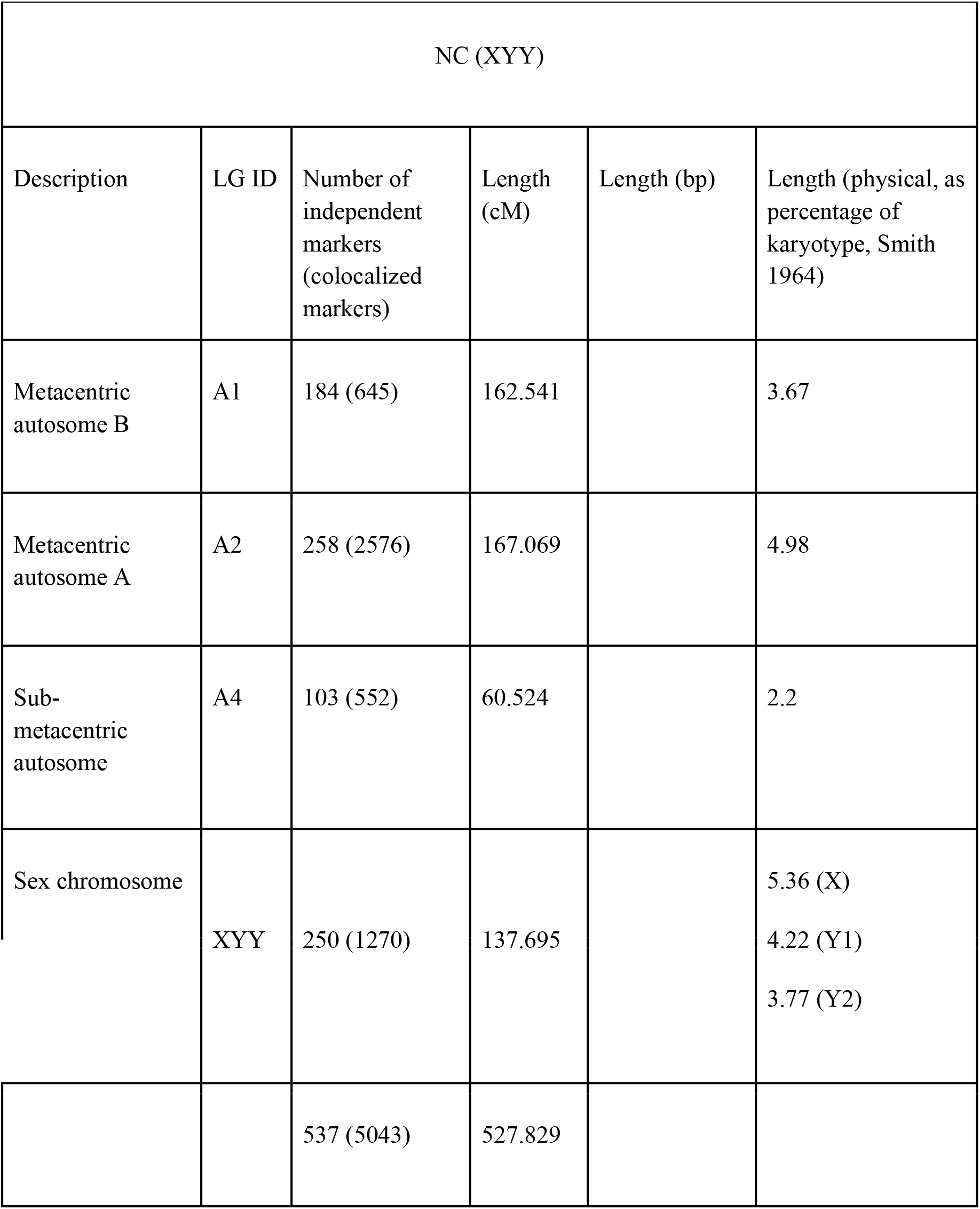
Major linkage groups in both maps of *Rumex hastatulus* LG ID numbers after Smith 1964

**Table S4.** Sex-linked SNPs of *Rumex hastatulus* based on Hough et al. (2014) pedigree data. TX = XY, NC = XYY.

|  |  |  |  |  |  |  |  |  |
| --- | --- | --- | --- | --- | --- | --- | --- | --- |
|  | Y-linked (TX) | Y-linked (NC) | X-linked (TX) | X-linked (NC) | Hemizygous (TX) | Hemizygous (NC) | Autosomal (TX) | Autosomal (NC) |
| A1 | 14 | 57 | 5 | 7 | 1 | 1 | 3894 | 1692 |
| A2 | 3 | 137 | 1 | 18 | 13 | 5 | 1700 | 3950 |
| A3 | 12 | 1172 | 11 | 763 | 6 | 7 | 1614 | 410 |
| A4 | 4 | 59 | 0 | 33 | 2 | 0 | 1173 | 1691 |
| X | 1208 | 980 | 439 | 456 | 50 | 79 | 420 | 348 |
| Not placed | 1048 | 1110 | 143 | 173 | 317 | 345 | 1681 | 1714 |

**Table S5.**
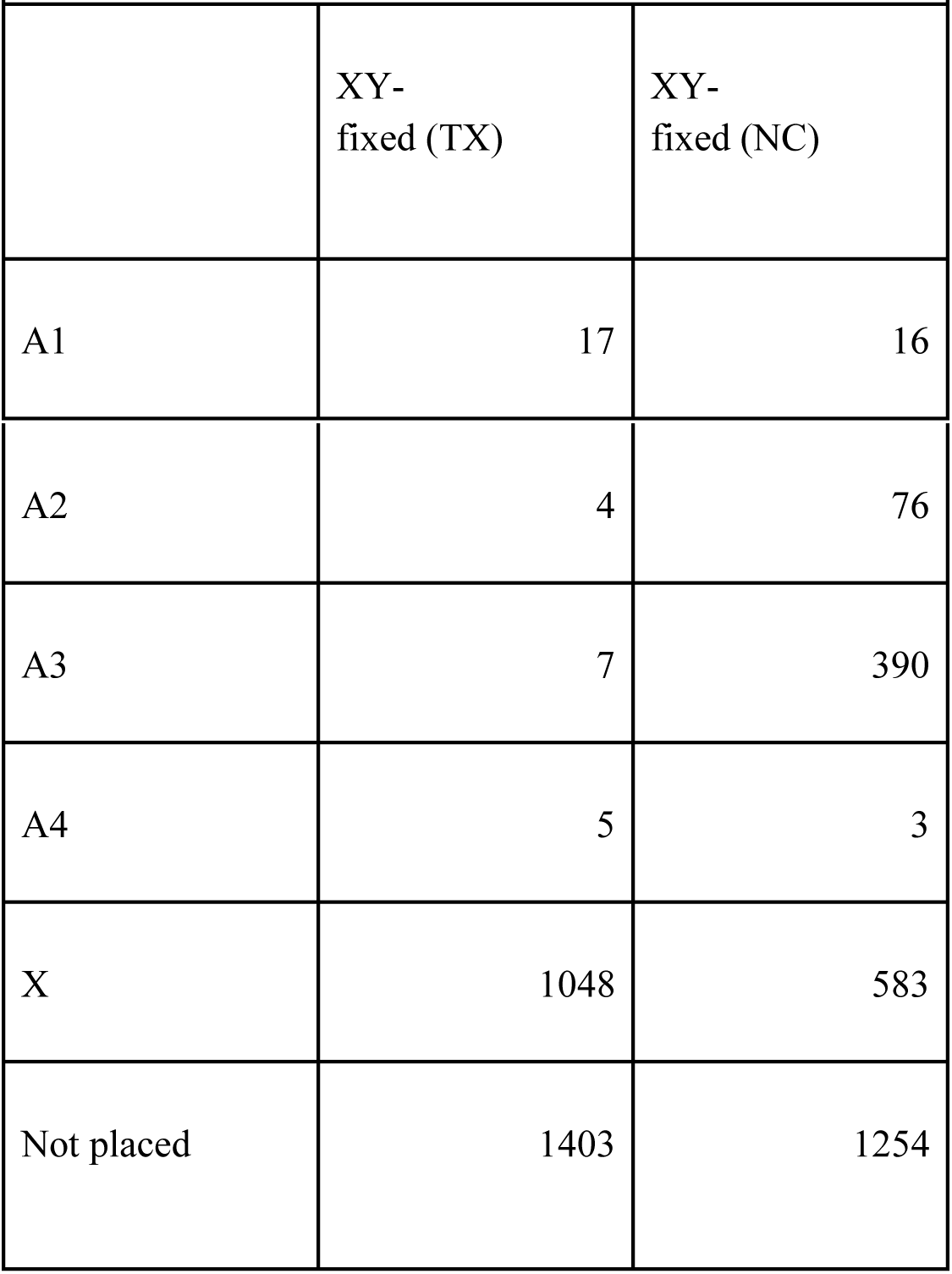
Sex-linked SNPs of *Rumex hastaulus* based on Hough et al. (2014) and Beaudry et al. (2019) population data. TX = XY, NC = XYY.

| **Table S5.** Sex-linked SNPs of *Rumex hastatulus* based on Hough et al. (2014) and Beaudry et al. (2019) population data. TX = XY, NC = XYY. |  |  |
| --- | --- | --- |
|  | XY-fixed (TX) | XY-fixed (NC) |
| A1 | 17 | 16 |
| A2 | 4 | 76 |
| A3 | 7 | 390 |
| A4 | 5 | 3 |
| X | 1048 | 583 |
| Not placed | 1403 | 1254 |

## Supplementary Figures

**Figure S1.**
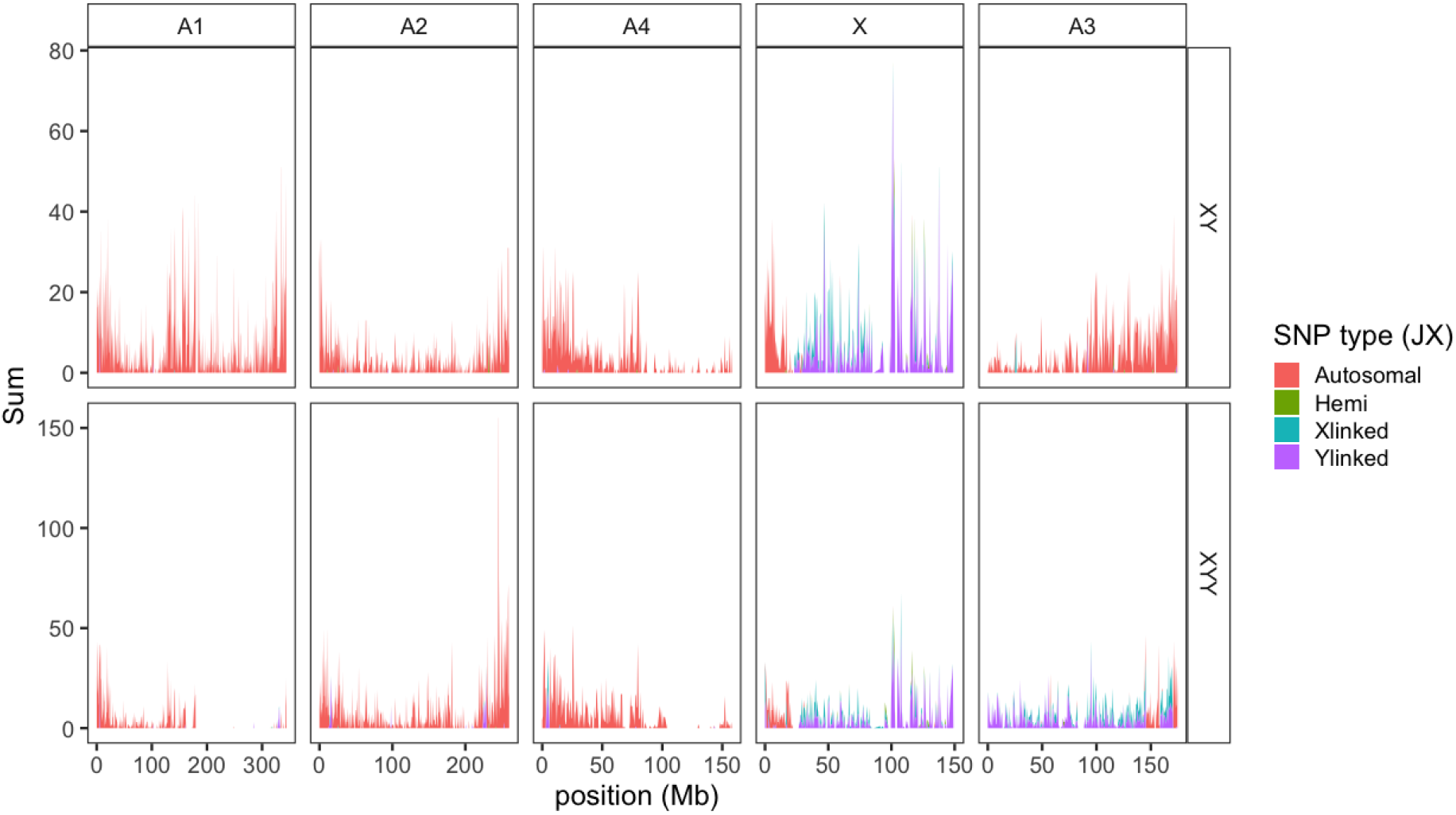
Windowed counts of SNPs identified as autosomal (red), hemizygous (green), and X-(blue) or Y-linked (purple) for the XY cytotype (top panel) and XYY cytotype (bottom-panel) of *Rumex hastatulus* in cross data from (35).

## References

1. B. S. Gaut, S. I. Wright, C. Rizzon, J. Dvorak, L. K. Anderson, Recombination: an underappreciated factor in the evolution of plant genomes. Nat. Rev. Genet. 8, 77–84 (2007).

2. M. Kirkpatrick, N. Barton, Chromosome inversions, local adaptation and speciation. Genetics 173, 419–434 (2006).

3. W. G. Hill, A. Robertson, The effect of linkage on limits to artificial selection. Genet. Res. 8, 269–294 (1966).

4. J. Felsenstein, The evolutionary advantage of recombination. Genetics 78, 737–756 (1974).

5. J. M. Smith, J. Haigh, The hitch-hiking effect of a favourable gene. Genet. Res. 23, 23–35 (1974).

6. B. Charlesworth, M. T. Morgan, D. Charlesworth, The effect of deleterious mutations on neutral molecular variation. Genetics 134, 1289–1303 (1993).

7. B. Charlesworth, D. Charlesworth, Selection of new inversions in multi-locus genetic systems. Genet. Res. 21, 167–183 (1973).

8. Q. Haenel, T. G. Laurentino, M. Roesti, D. Berner, Meta-analysis of chromosome-scale crossover rate variation in eukaryotes and its significance to evolutionary genomics. Mol. Ecol., 2477–2497 (2018).

9. K. Ohta, T. Shibata, A. Nicolas, Changes in chromatin structure at recombination initiation sites during yeast meiosis. EMBO J. 13, 5754–5763 (1994).

10. T. V. Kent, J. Uzunović, S. I. Wright, Coevolution between transposable elements and recombination. Philos. Trans. R. Soc. Lond. B Biol. Sci. 372 (2017).

11. L. H. Rieseberg, C. Van Fossen, A. M. Desrochers, Hybrid speciation accompanied by genomic reorganization in wild sunflowers. Nature 375, 313 (1995).

12. M. A. F. Noor, K. L. Grams, L. A. Bertucci, J. Reiland, Chromosomal inversions and the reproductive isolation of species. Proc. Natl. Acad. Sci. 98, 12084–12088 (2001).

13. D. Charlesworth, B. Charlesworth, Sex differences in fitness and selection for centric fusions between sex-chromosomes and autosomes. Genet. Res. 35, 205–214 (1980).

14. W. R. Rice, Sex chromosomes and the evolution of sexual dimorphism. Evolution 38, 735–742 (1984).

15. R. L. Trivers, “Parental investment and sexual selection” in Sexual Selection and the Descent of Man, 1871-1971, B. Campbell, Ed. (Aldine, 1972), pp. 136–179.

16. R. Lande, Sexual dimorphism, sexual selection, and adaptation in polygenic characters. Evolution 34, 292–305 (1980).

17. J. E. Mank, Sex chromosomes and the evolution of sexual dimorphism: lessons from the genome. Am. Nat. 173, 141–150 (2009).

18. W. R. Rice, The accumulation of sexually antagonistic genes as a selective agent promoting the evolution of reduced recombination between the primitive sex chromosomes. Evolution 41, 911–914 (1987).

19. L. H. Rieseberg, Chromosomal rearrangements and speciation. Trends Ecol. Evol. 16, 351–358 (2001).

20. J. J. Bull, Sex determining mechanisms: An evolutionary perspective. Experientia 41, 1285–1296 (1985).

21. J. F. Kidwell, M. T. Clegg, F. M. Stewart, T. Prout, Regions of stable equilibria for models of differential selection in the two sexes under random mating. Genetics 85, 171–183 (1977).

22. S. P. Otto, Evolutionary potential for genomic islands of sexual divergence on recombining sex chromosomes. New Phytol. (2019).

23. B. T. Lahn, D. C. Page, Four evolutionary strata on the human X chromosome. Science 286, 964–967 (1999).

24. L.-J. L. Handley, H. Ceplitis, H. Ellegren, Evolutionary strata on the chicken Z chromosome: implications for sex chromosome evolution. Genetics 167, 367–376 (2004).

25. R. Bergero, A. Forrest, E. Kamau, D. Charlesworth, Evolutionary strata on the X chromosomes of the dioecious plant *Silene latifolia*: evidence from new sex-linked genes. Genetics 175, 1945–1954 (2007).

26. R. Ming, A. Bendahmane, S. S. Renner, Sex chromosomes in land plants. Annu. Rev. Plant Biol. 62, 485–514 (2011).

27. D. Charlesworth, Plant sex chromosome evolution. J. Exp. Bot. 64, 405–420 (2013).

28. B. W. Smith, The evolving karyotype of *Rumex hastatulus*. Evolution 18, 93–104 (1964).

29. M. Kasjaniuk, A. Grabowska-Joachimiak, A. J. Joachimiak, Testing the translocation hypothesis and Haldane’s rule in *Rumex hastatulus*. Protoplasma 256, 237–247 (2019).

30. A. Grabowska-Joachimiak, et al., Chromosome landmarks and autosome-sex chromosome translocations in *Rumex hastatulus*, a plant with XX/XY1Y2 sex chromosome system. Chromosom. Res. 23, 187–197 (2015).

31. B. W. Smith, Evolution of sex-determining mechanisms in *Rumex*. Chromosom. Today 2, 172–182 (1969).

32. N. H. Putnam, et al., Chromosome-scale shotgun assembly using an in vitro method for long-range linkage. Genome Res. 26, 342–350 (2016).

33. E. Lieberman-Aiden, et al., Comprehensive mapping of long-range interactions reveals folding principles of the human genome. Science 326, 289–293 (2009).

34. J.-M. Belton, et al., Hi-C: a comprehensive technique to capture the conformation of genomes. Methods 58, 268–276 (2012).

35. J. Hough, J. D. Hollister, W. Wang, S. C. H. Barrett, S. I. Wright, Genetic degeneration of old and young Y chromosomes in the flowering plant *Rumex hastatulus*. Proc. Natl. Acad. Sci. 111, 7713–7718 (2014).

36. F. E. G. Beaudry, S. C. H. Barrett, S. I. Wright, Ancestral and neo-sex chromosomes contribute to population divergence in a dioecious plant. Evolution 54, 180 (2019).

37. M. P. H. Stumpf, G. A. T. McVean, Estimating recombination rates from population-genetic data. Nat. Rev. Genet. 4, 959–968 (2003).

38. A. Auton, G. McVean, Recombination rate estimation in the presence of hotspots. Genome Res. 17, 1219–1227 (2007).

39. M. M. Mahtani, H. F. Willard, Physical and genetic mapping of the human X chromosome centromere: repression of recombination. Genome Res. 8, 100–110 (1998).

40. D. Charlesworth, Young sex chromosomes in plants and animals. New Phytol. 224, 1095–1107 (2019).

41. D. Charlesworth, Does sexual dimorphism in plants promote sex chromosome evolution? Environ. Exp. Bot. 146, 5–12 (2018).

42. J. D. Fry, The genomic location of sexually antagonistic variation: some cautionary comments. Evolution 64, 1510–1516 (2010).

43. M. Iovene, Q. Yu, R. Ming, J. Jiang, Evidence for emergence of sex-determining gene(s) in a centromeric region in *Vasconcellea parviflora*. Genetics 199, 413–421 (2015).

44. S. M. Pilkington, et al., Genetic and cytological analyses reveal the recombination landscape of a partially differentiated plant sex chromosome in kiwifruit. BMC Plant Biol. 19, 172 (2019).

45. R. ten Hoopen, R. M. Harbord, T. Maes, N. Nanninga, T. P. Robbins, The self-incompatibility (*S*) locus in Petunia hybrida is located on chromosome III in a region, syntenic for the Solanaceae. Plant J. 16, 729–734 (1998).

46. S. P. Otto, B. A. Payseur, Crossover interference: shedding light on the evolution of recombination. Annu. Rev. Genet. 53, 19–44 (2019).

47. D. Berner, M. Roesti, Genomics of adaptive divergence with chromosome-scale heterogeneity in crossover rate. Mol. Ecol. 26, 6351–6369 (2017).

48. R. Navajas-Pérez, et al., The evolution of reproductive systems and sex-determining mechanisms within *Rumex* (Polygonaceae) inferred from nuclear and chloroplastidial sequence data. Mol. Biol. Evol. 22, 1929–1939 (2005).

49. J. Hough, W. Wang, S. C. H. Barrett, S. I. Wright, Hill-Robertson interference reduces genetic diversity. Genetics 207, 685–695 (2017).

50. G. Sandler, F. E. G. Beaudry, S. C. H. Barrett, S. I. Wright, The effects of haploid selection on Y chromosome evolution in two closely related dioecious plants. Evol. Lett. 197, 368–377 (2018).

51. M. Hartfield, S. P. Otto, P. D. Keightley, The role of advantageous mutations in enhancing the evolution of a recombination modifier. Genetics 184, 1153–1164 (2010).

52. N. Rodrigues, T. Studer, C. Dufresnes, N. Perrin, Sex-chromosome recombination in common frogs brings water to the fountain-of-youth. Mol. Biol. Evol. 35, 942–948 (2018).

53. D. Hojsgaard, E. Hörandl, A little bit of sex matters for genome evolution in asexual plants. Front. Plant Sci. 6, 82 (2015).

54. M. F. Scott, S. P. Otto, Haploid selection favors suppressed recombination between sex chromosomes despite causing biased sex ratios. Genetics 207, 1631–1649 (2017).

55. M. Pickup, S. C. H. Barrett, The influence of demography and local mating environment on sex ratios in a wind-pollinated dioecious plant. Ecol. Evol. 3, 629–639 (2013).

56. J. Eid, et al., Real-time DNA sequencing from single polymerase molecules. Science 323, 132–138 (2009).

57. C.-S. Chin, et al., Phased diploid genome assembly with single-molecule real-time sequencing. Nat. Methods 13, 1050–1054 (2016).

58. C.-S. Chin, et al., Nonhybrid, finished microbial genome assemblies from long-read {SMRT} sequencing data. Nat. Methods 10, 563–569 (2013).

59. A. Dobin, T. R. Gingeras, Mapping RNA-seq reads with STAR. Curr. Protoc. Bioinformatics 51, 11.14.1–11.14.9 (2015).

60. P. Danecek, et al., The variant call format and VCFtools. Bioinformatics 27, 2156–2158 (2011).

61. J. Taylor, D. Butler, R package ASMap: efficient genetic linkage map construction and diagnosis (2017).

62. Y. Wu, P. R. Bhat, T. J. Close, S. Lonardi, Efficient and accurate construction of genetic linkage maps from the minimum spanning tree of a graph. PLoS Genet. 4 (2008).

63. K. W. Broman, S. Sen, A Guide to QTL Mapping with R/qtl (2009) https:/doi.org/10.1007/978-0-387-92125-9.

64. A. Chakravarti, A graphical representation of genetic and physical maps: the Marey map. Genomics 11, 219–222 (1991).

65. F. E. G. Beaudry, S. C. H. Barrett, S. I. Wright, Genomic loss and silencing on the Y chromosomes of *Rumex*. Genome Biol. Evol. 9, 3345–3355 (2017).

66. F. J. Sedlazeck, P. Rescheneder, A. Von Haeseler, NextGenMap: Fast and accurate read mapping in highly polymorphic genomes. Bioinformatics 29, 2790–2791 (2013).

67. K. W. Broman, et al., R/qtl2: software for mapping quantitative trait loci with high-dimensional data and multiparent populations. Genetics 211, 495–502 (2019).

68. X. Zhou, M. Stephens, Genome-wide efficient mixed-model analysis for association studies. Nat. Genet. 44, 821–824 (2012).

